# A Virus-Packageable CRISPR Screen Identifies Host Factors Mediating Interferon Inhibition of HIV

**DOI:** 10.1101/363432

**Authors:** Molly Ohainle, Louisa Pendergast, Jolien Vermiere, Ferdinand Roesch, Daryl Humes, Ryan Basom, Jeffrey J. Delrow, Julie Overbaugh, Michael Emerman

## Abstract

Interferon (IFN) inhibits HIV replication by inducing an array of antiviral effectors. Here we describe a novel CRISPR knockout screening approach to identify the ensemble of these HIV restriction factors. We assembled a CRISPR sgRNA library specific for Interferon Stimulated Genes (ISGs) into a modified lentiviral vector that allows for packaging of sgRNA-encoding genomes *in trans* into budding HIV-1 particles. We observed that knockout of Zinc Antiviral Protein (ZAP) improved the performance of the screen due to ZAP-mediated inhibition of the vector. We identify a small panel of IFN-induced HIV restriction factors, including MxB, IFITM1, Tetherin/BST2 and TRIM5 which together explain the inhibitory effects of IFN on the HIV-1 LAI strain in THP-1 cells. Further, we identify novel HIV dependency factors, including SEC62 and TLR2. The ability of IFN-induced restriction factors to inhibit an HIV strain to replicate in human cells suggests that these human restriction factors are incompletely antagonized.

## Introduction

The HIV-1 pandemic resulted from a series of successive cross-species transmissions of primate lentiviruses. Simian Immunodeficiency Virus (SIV) transmission from African Old World primates to chimpanzees yielded the recombinant virus SIVcpz, which ultimately crossed into humans (Sharp & Hahn, 2011). Successful replication of lentiviruses in a new host species required adaptation to restriction factors in the new host (Etienne et al., 2015; Etienne, Hahn, Sharp, Matsen, & Emerman, 2013; Kirmaier et al., 2010). Restriction factors that target primate lentiviruses include TRIM5alpha, MxB, Tetherin, SAMHD1, the APOBEC3 family of cytidine deaminases (Malim & Bieniasz, 2012) and more recently described factors such as SERINC3/5, Zinc Antiviral Protein (ZAP), GBP5, SLFN11, LGALS3BP (90K), the HUSH complex, (Chougui et al., 2018; Krapp et al., 2016; M. Li et al., 2012; Lodermeyer et al., 2013; Rosa et al., 2015; Takata et al., 2017; Usami, Wu, & Gottlinger, 2015), as well as nearly 200 other proposed factors (reviewed in (Gelinas, Gill, & Hyde, 2018). HIV-1 has evolved accessory proteins that degrade many host restriction factors (Duggal & Emerman, 2012). Further, mutations preventing recognition by restriction factors, such as evolution of low CG dinucleotide content in the HIV-1 genome (Takata et al., 2017) or mutations in capsid (Kirmaier et al., 2010; F. Wu et al., 2013), represent another mechanism of escape.

Many restriction factors that target HIV-1 are induced by type I Interferon (IFN) and are therefore Interferon-Stimulated Genes (ISGs). Interferon has been implicated in at least partial control of HIV replication in chronically-infected individuals treated with IFN (Asmuth et al., 2010; Azzoni et al., 2013) as well as in SIV-infected rhesus macaques (Sandler et al., 2014). In contrast, IFN levels have also been correlated with higher viral load and decreased CD4 T cell counts in HIV-infected individuals (Hardy et al., 2013). Further, it appears that ISG expression exerts changing selective pressure on HIV evolution *in vivo* since transmitted/founder (T/F) strains are relatively resistant to IFN compared to viruses isolated later in infection (Fenton-May et al., 2013; Iyer et al., 2017; Parrish et al., 2013). It remains to be determined if one dominant ISG mediates all or most of the IFN inhibition, or if a multitude of antiviral ISGs together limit viral replication in response to IFN.

The HIV-1 LAI strain (HIV-1_LAI_) was isolated from a chronically-infected individual (Wain-Hobson et al., 1991) and is sensitive to type I IFN. Specifically, potent IFNα inhibition of HIV-1_LAI_ can be observed in the THP-1 monocytic cell line (Goujon & Malim, 2010). MxB, an interferon-induced GTPase that binds to and blocks lentiviral capsids, was identified as an IFNα-induced factor in THP-1 cells (Goujon et al., 2013; Kane et al., 2013; Z. Liu et al., 2013), although the role of MxB in the IFNα-induced inhibition of HIV infection in these cells has been questioned (Opp, Vieira, Schulte, Chanda, & Diaz-Griffero, 2015). Restriction factors have previously been discovered through cDNA library screening or by comparing expression of transcript levels in permissive versus non-permissive cells (Goujon et al., 2013; Kane et al., 2013; Neil, Zang, & Bieniasz, 2008; Sheehy, Gaddis, Choi, & Malim, 2002; Stremlau et al., 2004). More high-throughput approaches to find HIV restriction factors have focused on either overexpression screens to identify broad antiviral ISGs (Schoggins et al., 2011) or HIV-specific antiviral ISGs (Kane et al., 2016). Further, one screen for HIV restriction factors was also performed by transfection of siRNA pools (L. Liu et al., 2011). However, a more complete understanding of the constellation of restriction factors that inhibit HIV in human cells and a more tractable, high-throughput method to discover restriction factors remains to be described.

Here we describe a CRISPR/Cas9-mediated gene knockout functional screening method in which lentiviral genomes encoding CRISPR sgRNAs are packaged into budding HIV virions, allowing robust identification of HIV restriction factors and dependency factors in a high-throughput manner. Cas9 endonuclease and sgRNA are delivered to cells in a vector that is modified to be transcribed and subsequently packaged *in trans* by the infecting HIV virus. Deep sequencing of packaged HIV-CRISPR RNA in nascent HIV virions released from pooled KO cells serves to proxy the efficiency of HIV replication in each genetic knockout. Thereby, our approach allows for targeted gene knockout and a functional assay simultaneously across thousands of genes in a heterogeneous population of cells, i.e., multiplexed host factor screening. Furthermore, as read-out of the functional assay is done of at the level of newly budded viruses, the approach allows for screening of restrictions factors affecting the full HIV life cycle. We find a small panel of ISGs to mediate IFN inhibition of HIV-1 in THP-1 cells, including MxB, TRIM5alplha, IFITM1 and Tetherin. Further, this approach can as be used to identify HIV dependency factors and we identify CD169, SEC62 and TLR2 as important host factors in THP-1 cells. The results presented here suggest that adaptation of primate lentiviruses to humans is incomplete as we find that the same host restriction factors that block cross-species transmission also play a role in limiting the replication of highly-adapted HIV-1 in IFN-stimulated cells.

## Results

### An ISG-specific knockout screen that packages sgRNA-encoding lentiviral genomes into virions

IFNα inhibits HIV_LAI_ replication in THP-1 cells 10-fold (Goujon et al., 2013). To identify the factor(s) mediating the IFNα-induced inhibition of HIV, we designed a novel HIV-based CRISPR screen in which the virus itself serves as a reporter. Cells which lack a dependency factor due to CRISPR-mediated gene knockout will release less virus, whereas cells which lack a restriction factor will produce more virus as compared to control cells which containing single-guide RNA (sgRNA) sequences that do not target any human genes, Non-Targeting Controls (NTCs). We engineered a Cas9 and sgRNA-encoding lentiviral vector such that sgRNA-encoding genomic RNA can be packaged *in trans* by budding HIV virions. Therefore, the normalized abundance of Cas9/sgRNA-encoding genomes themselves are the direct readout for the functional activity of each gene knockout on viral replication. Importantly, this approach will allow for assay of effects of gene knockout on a complete round of viral replication. The lentiCRISPRv2 lentiviral vector contains a Self-Inactivating (SIN) LTR that prevents transcription after integration (Shalem, Sanjana, Hartenian, Shi, Scott, Mikkelsen, et al., 2014). We repaired this 3’ LTR with a complete HIV-1 LTR, creating a transcription- and packaging-competent construct we call HIV-CRISPR (Figure 1A).

**FIGURE 1.**
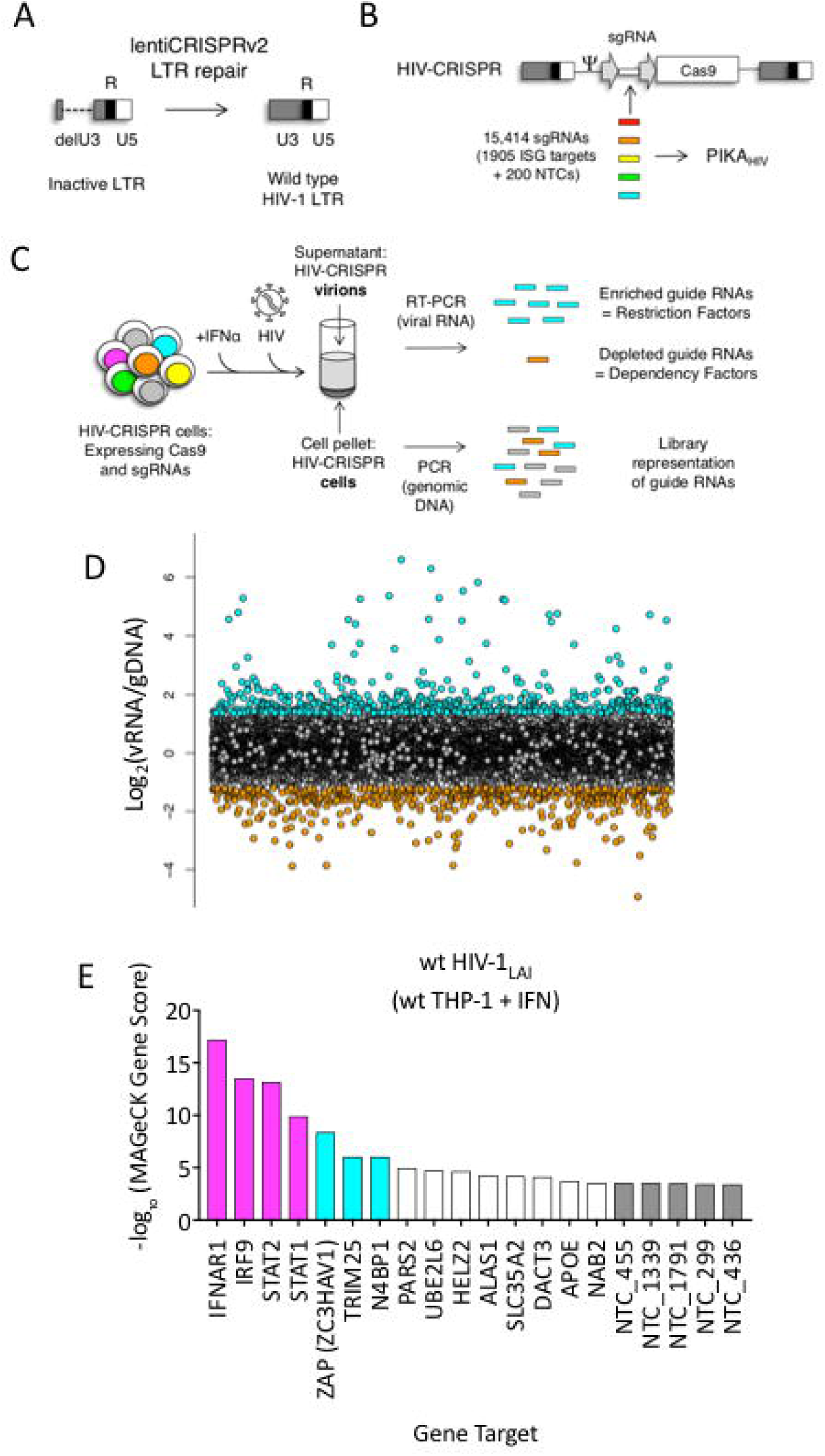
The HIV-CRISPR screen identifies gene knockouts that increase and decrease HIV infection. <u>A</u>: The deletion in the U3 region of the SIN LTR vector (lentiCRISPRv2) was repaired by inserting the full-length HIV-1_LAI_ LTR sequence to create the HIV-CRISPR construct. <u>B</u>: The ISG-targeting sgRNA library (15,348 unique sgRNA sequences) was synthesized and assembled into the HIV-CRISPR backbone to create the PIKA_HIV_ (<u>P</u>ackageable <u>I</u>SG <u>K</u>nockout <u>A</u>ssembly) library. <u>C</u>: THP-1 cells containing the PIKA_HIV_ CRISPR knockout library were stimulated overnight with 1000 U/mL IFNα and infected with HIV-1_LAI_ in duplicate infections. Viral RNA and genomic DNA were collected 3 days post infection and sgRNA sequences present in virions (vRNA) and genomic DNA (gDNA) were quantified through RT-PCR/PCR and deep sequencing. <u>D</u>: Average enrichment of sgRNA sequences in the viral RNA (vRNA) was compared to their representation in the sequenced genomic DNA (gDNA) (n=2). <u>Y-Axis</u>: Log_2_-normalized Fold Change (log_2_FC) of vRNA sgRNA sequences as compared to gDNA sgRNA sequences. <u>X-Axis</u>: random order of individual sgRNAs. The 200 non-targeting control (NTC) sgRNAs are shown in gray. The most enriched sgRNA sequences are in cyan (top 500), most depleted in orange (bottom 500) and other sgRNA sequences are in white. <u>E</u>: MAGeCK Gene analysis was performed to identify the highest-scoring genes based on sgRNA frequencies for each gene across both replicates. An NTC gene set was created *in silico* by iteratively binning the 200 Non-Targeting Control sgRNA sequences into NTC Genes. <u>Y-Axis</u>: −log_10_MAGeCK Gene Score. The type I IFN pathway genes IFNAR1, IRF9, STAT1 and STAT2 are shown in magenta. Non-Targeting Controls (NTCs) are in gray. Candidate Hits are cyan. The top 20-scoring genes across replicate screens are shown.

To target genes mediating the IFN inhibition of HIV-1, we curated a list of potential ISGs from existing microarray and RNA-seq datasets from cell types relevant to HIV-1 infection, including PBMCs, primary CD4+ T cells, monocyte-derived macrophages (MDMs), monocytes and the THP-1 monocytic cell line (Supplemental Figure S1A and Supplemental Table S1). Thus, the library is also enriched in genes that are specifically expressed in HIV target cells. For each of the 1905 ISGs present in our library, we selected a total of 8 sgRNA sequences from existing human whole-genome CRISPR/Cas9 libraries (Supplemental Figure S1B and Supplementary Table S2) (Doench et al., 2016; Hart et al., 2015; Sanjana, Shalem, & Zhang, 2014b; Shalem, Sanjana, Hartenian, Shi, Scott, Mikkelson, et al., 2014; Shalem, Sanjana, & Zhang, 2015; T. Wang et al., 2015; T. Wang, Wei, Sabatini, & Lander, 2014). 200 Non-Targeting Control (NTC) sgRNA sequences that are not predicted to target any loci in the human genome were also included (Shalem et al., 2015) (Supplemental Figure S1B and Supplementary Table S2). In total 15,348 unique sgRNA sequences were assembled into the HIV-CRISPR backbone to create the Packageable ISG Knockout Assembly or PIKA_HIV_ library (Figure 1B). The enrichment or depletion of sgRNA sequences in the viral RNA (vRNA) as compared to the genomic DNA (gDNA) of the cells is quantified through sequencing of sgRNA sequences both in released HIV particles and integrated into the cellular genomic DNA. sgRNAs that target antiviral genes (restriction factors) are overrepresented in viral supernatants due to more robust viral replication specifically in these KO cells (Figure 1C, cyan). Conversely, sgRNAs that target dependency factors are depleted in viral supernatants due to decreased viral replication specifically in these KO cells (Figure 1C, orange).

To perform the screen, 8 × 10^6^ THP-1 cells were transduced with the PIKA_HIV_ library at an MOI < 1 (MOI = 0.6) to create a population of cells with single HIV-CRISPR integrations at >500X coverage. THP-1/PIKA_HIV_ cells were split in two independent replicates and left untreated or treated with IFNα overnight. Each replicate was then infected with HIV-1 at a dose that infects 50% of cells without IFNα treatment. Secreted virus was collected 3 days after infection, and sgRNA sequences encoded by HIV-CRISPR genomic RNA packaged into budding HIV virions were amplified by RT-PCR and quantitated through deep sequencing (Figure 1C). THP-1/PIKA_HIV_ cells were collected in parallel at the time of viral supernatant harvest and the genomic DNA (gDNA) was also deep sequenced. We compared the relative enrichment of HIV-CRISPR sgRNA sequences in the viral RNA (vRNA) to the genomic DNA (gDNA) to find enriched and depleted sgRNA sequences (Figure 1D and Supplemental Table S3). Relative to the NTCs (Figure 1D, gray circles), there are a number of sgRNA sequences that are either enriched (Figure 1D, top 500 sgRNAs in cyan) or depleted (Figure 1D, bottom 500 sgRNAs in orange) in the viral supernatant as compared to the NTCs. Since each gene in the PIKA_HIV_ library is targeted by 8 individual sgRNAs, we analyzed the enrichment across all sgRNAs for a gene using the MAGeCK package across both duplicates (W. Li et al., 2014). We identify the type I IFN pathway genes, STAT1, IFNAR1, STAT2 and IRF9 as the highest-scoring hits (magenta in Figure 1E). Therefore, the PIKA_HIV_ screen functions as designed: cells in which IFN signaling is compromised exhibit increased viral production and, therefore, enriched HIV-CRISPR representation of sgRNAs in the secreted HIV virions. After the IFN pathway genes, the Zinc Antiviral Protein (ZAP) and its modifier TRIM25 were the next to highest scoring hits. ZAP is an antiviral effector that has potent activity against alphaviruses as well as moderate activity against retroviruses (Bick et al., 2003; Gao, Guo, & Goff, 2002; Kerns, Emerman, & Malik, 2008; Takata et al., 2017). TRIM25 is a gene known to modify ZAP’s antiviral activity (M. M. Li et al., 2017). More recently, it was shown that ZAP blocks virus replication by degrading transcripts with a high CG dinucleotide content (Takata et al., 2017). We also find NEDD4 Binding Protein 1 (N4BP1), a poorly-characterized inhibitor of the E3 ligase ITCH in mice (Oberst et al., 2007) that has not been previously known for antiretroviral activity (Figure 1E; cyan). N4BP1 encodes RNA binding domains and is proposed to have RNase activity (Anantharaman & Aravind, 2006).

### An iterative PIKA_HIV_ screen in ZAP-KO cells identifies a panel of ISGs that inhibit HIV in THP-1 cells

We generated ZAP and N4BP1 knockout (KO) cell lines by electroporating crRNA/Cas9 complexes (crRNPs) into THP-1 cells and single-cell cloning and found that knockout of either gene only very modestly increases infection of HIV (Supplemental Figure S2A and S2B). Therefore, we reasoned that the true IFN-induced restriction factors that potently inhibit HIV in THP-1 cells were not identified in this initial screen. Analysis of the CG dinucleotide content across the HIV-CRISPR genome shows high levels of CG dinucleotides, particularly in the Cas9 and Puromycin resistance ORFs, that are potential targets for ZAP-mediated RNA degradation (Figure 2A). Given its role in degradation of RNA with high CG content, we hypothesized that ZAP could inhibit the full-length HIV-CRISPR genomic RNA that is packaged into budding virions rather than the wt HIV genome. Thus, we determined whether or not ZAP KO allows for increased packaging of the HIV-CRISPR vector in viral particles released from cells by measuring both wild type HIV-1_LAI_ genomes (HIV-Pol; black in Figure 2B) and HIV-CRISPR genomes (cPPT-U6; gray in Figure 2B) with a ddPCR assay. Indeed, we find enhanced packaging of HIV-CRISPR genomes relative to wild type HIV-1_LAI_ genomes in the viral supernatant in cell populations with reduced ZAP expression (Figure 2B - 10.5% in wt THP-1 cells; 24.8% and 31.6% in the ZAP-KO clonal lines).

**FIGURE 2.**
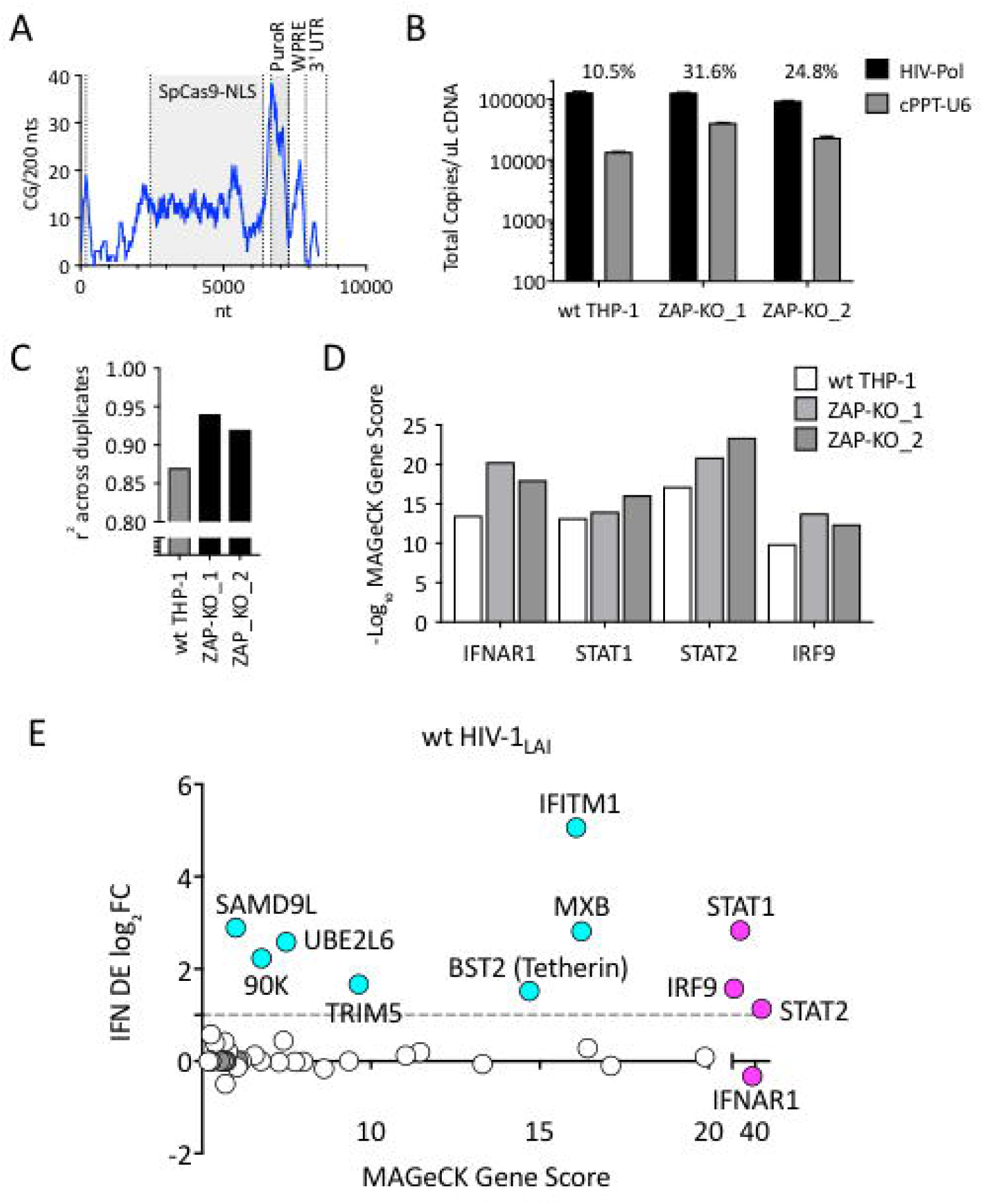
Iterative screening in ZAP-KO THP-1 cells Identifies IFNα-induced Inhibitors of HIV Replication. <u>A</u>: Sliding window analysis of CG dinucleotide content per 200 nucleotides of the HIV-CRISPR construct (blue line). The Cas9 and Puromycin coding regions are shaded gray. <u>B</u>: Total copies of wt HIV genomic RNA (HIV-Pol – black bars) and HIV-CRISPR genomic RNA (cPPT-U6 – gray bars) released from wild type THP-1 cells as compared to 2 clonal ZAP-KO THP-1 lines (ZAP-KO_1 and ZAP-KO_2) were assayed by ddPCR using primers specific for each vector template. ddPCR on cDNA from each infection was performed in duplicate. <u>C</u>: PIKA_HIV_ screen in IFNα-treated THP-1 cells with and without endogenous ZAP expression. Data for all sgRNAs in duplicates were compared to generate an overall r^2^ value. Correlation of read counts across duplicate screens in each cell type (gray = wild type THP-1 cells; black = ZAP-KO THP clonal lines). <u>D</u>: MAGeCK Gene Scores for the type I IFN pathway genes (IFNAR1, STAT1, STAT2, IRF9) across screens in wild type THP-1 (white bars) and in two ZAP-KO clonal THP-1 lines (gray bars). <u>E</u>: MAGeCK Gene Scores for Hits Identified in the THP-1 ZAP-KO cells. Positive MAGeCK Gene Scores for the results from both ZAP-KO screens was multiplied to generate a ZAP-KO MAGeCK Gene Score. <u>Y-Axis</u>: IFN induction (log2FoldChange) in THP-1 cells calculated from GSE46599. <u>X-Axis</u>: Combined MAGeCK Gene Scores for top 40 Hits in both ZAP-KO screens. Magenta: IFN pathway genes (IFNAR1, STAT1, STAT2, IRF9). Cyan: highly-IFN induced, high-scoring candidate Hits. White: other high-scoring hits which were not IFN-induced (full list in Supplemental Table S4).

Therefore, to circumvent the inhibitory effects of ZAP on the HIV-CRISPR vector, we repeated the PIKA_HIV_ screen in two ZAP-KO THP-1 clonal cell lines. As expected for a screen in ZAP-KO cells, ZAP is no longer a significantly-scoring hit in the screen (rank # 1647/3812 in combined ZAP-KO screen data; Supplemental Table S4). In addition, there is also no enrichment of N4BP1 or TRIM25 in the ZAP-KO screens (rank # 3789/3812 and 3090/3812 in combined ZAP-KO screen data; Supplemental Table S4), suggesting that the inhibitory activity of N4BP1 and TRIM25 in the HIV-CRISPR screen are ZAP-dependent.

To ask if ZAP knockout improves performance of the HIV-CRISPR screen, we analyzed read counts across duplicates in the two independent ZAP-KO THP-1 clonal lines and compared the results to the screen performed in wild-type THP-1 cells. There is better correlation in sgRNA representation across replicates performed in ZAP-KO THP-1 cells as compared to control THP-1 cells (Figure 2C; r^2^ = 0.92 and 0.94 for the ZAP-KO screens as compared to r^2^ = 0.87 for the screen in wild type THP-1 cells). Further, an analysis specifically across the four genes that are well-described components of the type I IFN pathway also show increased Gene Scores in the ZAP-KO THP-1 clonal lines (Figure 2D and Supplemental Table S4) suggesting that deletion of ZAP-mediated inhibition from THP-1 cells improves performance of the HIV-CRISPR screen.

By multiplying gene scores from both ZAP-KO screens (Figure 2E; MAGeCK score on x-axis) we identify a list of candidate hits. To ask which genes are most likely to contribute specifically to the IFN-mediated inhibition of HIV-1, we calculated the level of IFN induction of each of the top hits from an existing THP-1 microarray dataset (Figure 2E; IFN log_2_FC on y-axis and Supplemental Table S4). No hit scored as highly as the type I IFN pathway genes (magenta in Figure 2E). Therefore, multiple genes, rather than a single ISG, are responsible for the IFN-mediated inhibition of HIV infection in THP-1 cells. Further, a small subset of genes, including MxB, IFITM1, Tetherin, TRIM5, UBE2L6, LGALS3BP (90K) and SAMD9L, are candidate restriction factors mediating the IFN inhibition of HIV-1 in THP-1 cells. MxB, IFITM1, Tetherin and TRIM5alpha are the most significantly-scoring hits in the PIKA_HIV_ screen that are also highly-induced by IFN (Figure 2E). All have well-described anti-lentiviral functions (Goujon et al., 2013; Kane et al., 2013; Z. Liu et al., 2013; Lu et al., 2011; Malim & Bieniasz, 2012; Neil et al., 2008; Stremlau et al., 2004). Thus, the PIKA_HIV_ screen identifies IFN-induced restriction factors in a massively-parallel approach assaying all gene targets simultaneously in pools of knockout cells.

### MxB is a dominant mediator of the IFN inhibition of HIV-1 in THP-1 cells but its activity depends on the route of viral entry

To determine the relative importance of MxB to the IFN-induced block to infection, we created MxB KO THP-1 cells. MxB was deleted from THP-1 cells by transduction with a lentiCRISPRv2 MxB-targeting construct followed by single-cell cloning. Deletion of MxB expression was confirmed through western blot of IFN-treated clonal MxB-KO lines (Figure 3A). On creating clonal populations of THP-1 cells, we observed substantial heterogeneity across clonal lines of THP-1 cells (compare infection levels in NTC clonal lines in Figure 3B). Therefore, we infected many clonal NTC and MxB-KO cell lines in parallel. Infection of MxB-KO cells confirms that MxB plays a major role in the IFN block to infection as there is rescue of the IFN effect as compared to controls (Figure 3B and 3C; the Fold Inhibition in MxB-KO cells is close to 1). Therefore, MxB is a the dominant, early-acting ISG inhibiting HIV replication in THP-1 cells.

**FIGURE 3.**
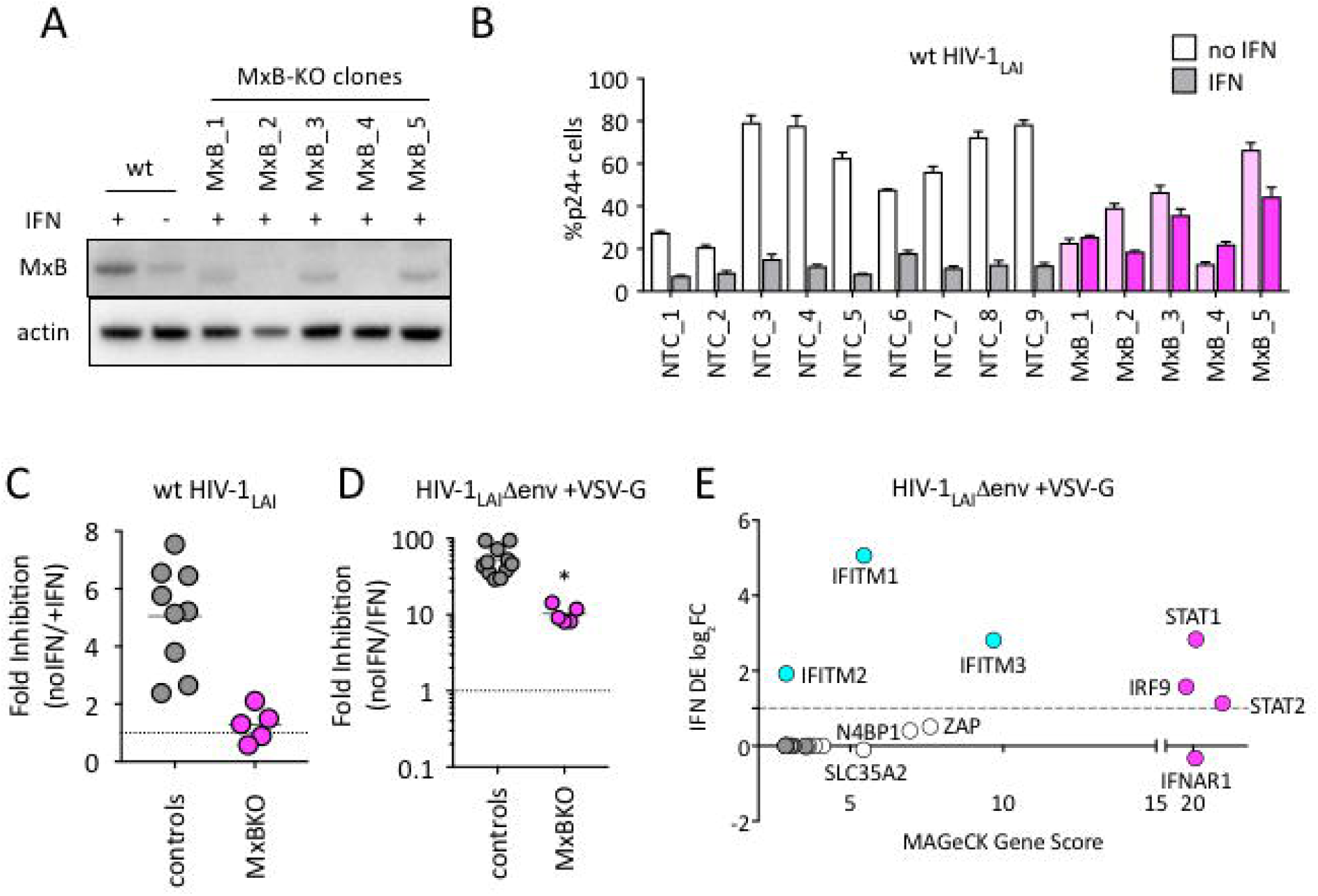
MxB is a dominant, early-acting ISG whose activity is masked by other ISGs when HIV entry is mediated by VSV-G. <u>A</u>: Clonal MxB-KO THP-1 lines generated by transducing with an MxB-targeting sgRNA/lentiCRISPRv2 construct, selection and single-cell sorting. Western blot for MxB expression with and without IFNα stimulation overnight is shown for wild type (wt) THP-1 cells; MxB-KO clones were all IFNα treated overnight. Note: the lower molecular weight band in some lanes results from initiation at an internal Met codon that would not be predicted to have anti-HIV activity (Goujon, Greenbury, Papaioannou, Doyle, & Malim, 2015; Matreyek et al., 2014). <u>B</u>: Nine individual clonal THP-1 lines (white/gray bars) along with 5 clonal THP-1 MxB-KO lines (pink bars) were pre-treated with IFNα overnight and infected with wt HIV-1_LAI_. The percentage of cells expressing HIV p24gag was assayed 2 days post-infection by intracellular staining and flow cytometry (n = 3). Light bars = no IFN; Dark bars = overnight IFNα treatment prior to infection. <u>C</u>: The Fold Inhibition (%p24+ cells without IFN/%p24+ cells with IFNα) calculated for each clonal line for wt HIV-1_LAI_ infections from the data in Panel B. Controls = gray; MxB-KO = magenta. Dotted line at a Fold Inhibition of 1 = no IFN inhibition. <u>D</u>: Individual clonal control THP-1 lines (gray) along with MxB-KO clonal lines (magenta) were infected with VSV-G pseudotyped HIV both with and without IFNα pretreatment (n = 3). Fold Inhibition was calculated as in C. Dotted line: Fold Inhibition of 1 = no IFN inhibition. *p=0.0014 (unpaired t test). <u>E</u>: The PIKA_HIV_ screen was performed in triplicate in wild type THP-1 cells. Y-Axis: IFN induction as determined by Differential Expression Analysis of microarray data in THP-1 cells (log_2_FoldChange). X-Axis: MAGeCK Gene Scores for Top 25 Hits. Magenta: IFN pathway genes (IFNAR1, STAT1, STAT2, IRF9). Cyan: highly-IFN induced, high-scoring candidate Hits. White: non-IFN induced genes including ZAP, N4BP1, and SLC35A2.

Viral entry is a key target of potent IFN-mediated restriction, specifically by ISGs such as IFITMs, a family of 5 membrane-resident antiviral genes in humans with broad antiviral effects (Shi, Schwartz, & Compton, 2017). IFITMs restrict viruses that enter cells by fusion at the plasma membrane or in the endosome. We hypothesized that sensitivity to MxB restriction may be dependent on the viral envelope since our previous work has shown that restriction of lentiviruses using distinct entry pathways are differentially affected by ISGs (Roesch, OhAinle, & Emerman, 2018). We found that while the IFN inhibition in the MxB-KO clonal lines is significantly lower than that of control clonal lines (Figure 3D; p = 0.014 unpaired t test), there is still a large inhibition of replication of VSV-G pseudotyped HIV-1 by IFNα (Figure 3D; 53-Fold). Thus, one or more ISGs induced by IFNα potently block VSV-G mediated entry in THP-1 cells independent of MxB. To ask what factors mediate this block, we repeated the HIV-CRISPR screen with VSV-G pseudotyped HIV-1 in THP-1 cells. In addition to ZAP and N4BP1, shared in common with the original screen (Figure 1), the antiviral proteins IFITM1, IFITM2 and IFITM3 are the most significantly-scoring hits (Figure 3E). This suggests that IFITMs are the dominant IFN-induced blocks to replication when HIV-1 is pseudotyped with the VSV-G envelope. Significant overlap in sequence across IFITM orthologues complicates interpretation of the screen data in terms of which IFITMs are most important, as some sgRNAs in our library likely target multiple IFITM loci. However, these results show that while MxB does play a role in the IFN-mediated inhibition of VSV-G pseudotyped HIV-1 viruses, this effect is masked by dominant IFITM inhibition of these pseudotyped viruses. Therefore, viral entry route impacts restriction factor sensitivity.

### TRIM5alpha, IFITM1 and Tetherin are additional ISGs that contribute to the IFN block

We were surprised to find TRIM5 and Tetherin in this screen as HIV-1 is thought to be highly-adapted to these human restriction factors. To assay the contribution of each of these ISGs to IFN inhibition of HIV in THP-1 cells, we measured viral replication in THP-1 KO pools. Pretreating cells with IFNα shows ~7-fold inhibition of infection in the control NTC cell pools (Figure 4A) while IFN-mediated inhibition of HIV was significantly lower in MxB, TRIM5 and IFITM1 KO lines than in NTCs pools (MxB_1 = 2.6-fold, MxB_2 = 2.5-fold, TRIM5_1 = 3.9-fold, TRIM5_2 = 4.8-fold, IFITM1_1 = 4.7-fold, IFITM1_2 = 6-fold and IFITM1_3 = 4.3-fold in Figure 4B; p<0.05). The largest rescue we observed was in the MxB knockout pools (Figure 4B), confirming that key role of MxB in the IFN phenotype. However, TRIM5 and IFITM1 also contribute to IFNα inhibition (Figure 4B). We find no effect of Tetherin KO on early steps of HIV replication as expected given its role as a late-acting HIV restriction factor (Tetherin_1 = 6-fold, Tetherin_2 = 6.4-fold in Figure 4B) (Neil et al., 2008; Van Damme et al., 2008).

**FIGURE 4.**
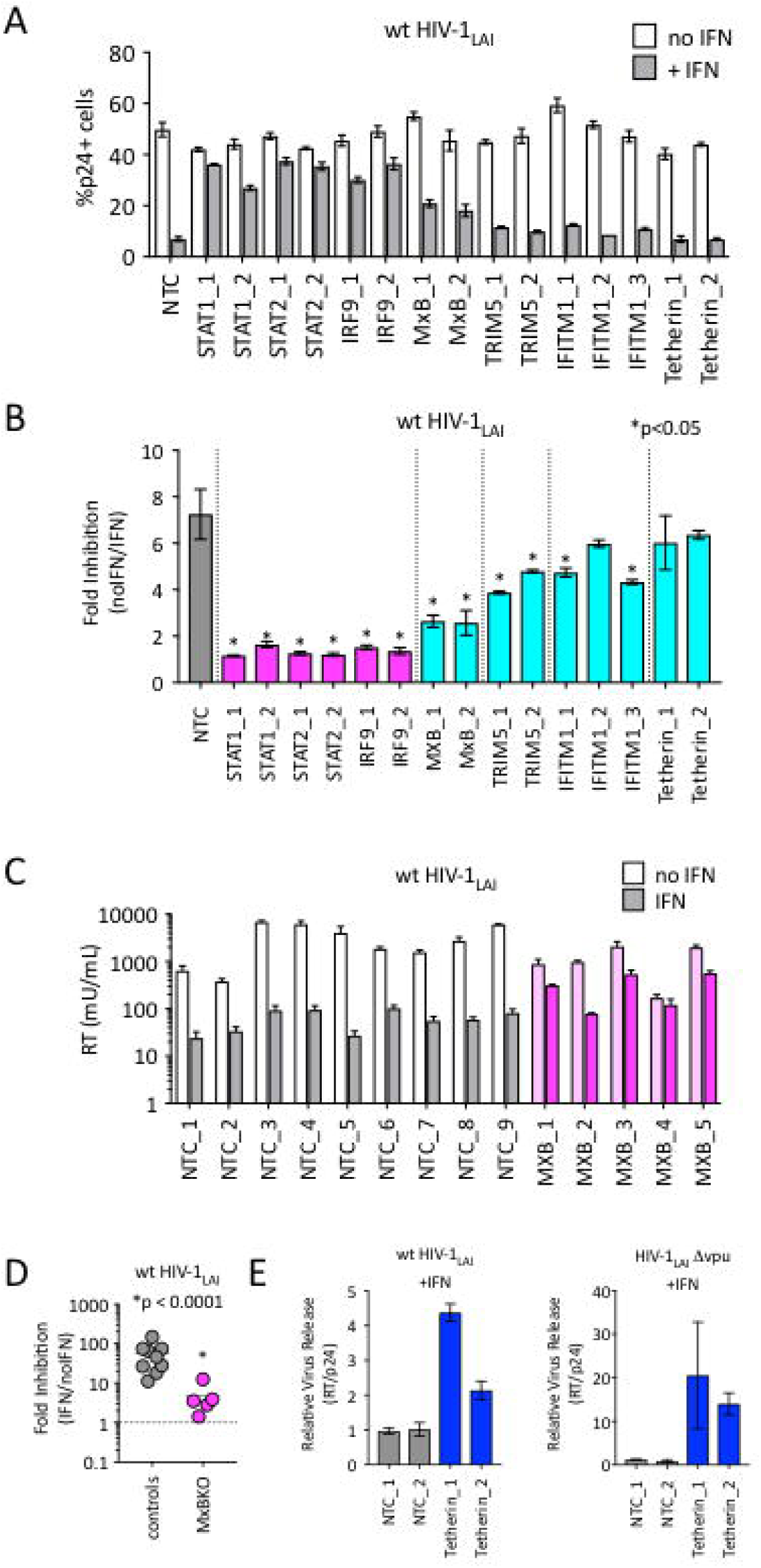
TRIM5alpha, IFITM1 and Tetherin are additional ISGs that contribute to the IFN block. <u>A</u>: THP-1 cell pools edited for gene targets of interest were created by transducing wild type THP-1 cells with lentiCRISPRv2 sgRNA constructs (2 different sgRNAs - for STAT1, STAT2, IRF9, MxB, TRIM5 and Tetherin, and 3 for IFITM1), selected for 2 weeks to allow gene knockout (see Supplemental Table 6 for analysis of gene knockout efficiency) and infected with HIV-1_LAI_ with and without IFNα pretreatment in triplicate (white bars = no IFNα, gray bars = +IFNα). The percentage of cells expressing HIV p24gag was assayed 2 days post-infection by intracellular staining and flow cytometry. NTC n=9. MxB_2 n=6. All other pools n=3. <u>B</u>: The Fold Inhibition (%p24+ cells without IFN/%p24+ cells with IFNα) is shown for each KO pool. Control (NTC) = gray; IFN pathway genes = magenta. MxB = Cyan. TRIM5 = Green. IFITM1 = Purple. Tetherin = Yellow. Cells pools with significantly reduced Fold Inhibition as compared to the NTC pools *p<0.05 (unpaired t test). <u>C</u>: Virus release from the clonal NTC (white/gray) or MxB-KO clones (pink) from Figure 3 as measured with a viral RT assay at 3 days post-infection with HIV-1_LAI_ with and without IFNα (light bars = no IFN; dark bars = IFNα). <u>D</u>: The Fold Inhibition (RT mU/mL without IFN/RT mU/mL with IFNα) calculated for each clonal line. Controls = gray; MxB-KO = magenta. Dotted line: Fold Inhibition of 1 = no IFN inhibition. <u>E</u>: THP-1 cell pools (NTC_1 and NTC_2 = gray; Tetherin-KO pools = blue) created by transduction with lentiCRISPRv2 lentiviral vectors were infected in triplicate with Vpu-deficient HIV (HIV-1_LAI_Δvpu) or wild type HIV (wt HIV-1_LAI_). The HIV_LAI_Δvpu contains a frameshift mutation in Vpu upstream of the env open reading frame. IFNα was added 16 hours post-infection and T-20 fusion inhibitor was added 24 hours post-infection. The amount of reverse transcriptase (RT) activity released into the supernatant was then normalized to the percentage of p24+ cells in order to directly quantify virus release per infected cell in the presence of IFN (RT mU/mL/%p24+ cells in culture).

We also observed a significant IFNα-mediated block to the late stages of the HIV lifecycle (after translation of the viral Gag protein used to detect infection in Figure 4A) in both control and MxB-KO cells (Figure 4C). While MxB-KO clonal lines show a decreased IFN effect compared to NTC clonal lines (compare controls to MxB-KOs in Figure 4D), there is still a 4.8-fold inhibition of virus released from MxB-KO clonal lines (magenta in Figure 4D). Since Tetherin is a well-characterized late-acting restriction factor and was also a hit in our PIKA_HIV_ screen, we asked if Tetherin is responsible for the late ISG block we observed. We assayed virus release from KO cell pools (NTC control, IFN Pathway genes: IFNAR1, STAT1, STAT2, IRF9 and Tetherin) when IFN was added 16 hours after infection to bypass early-acting ISGs. Infection with Vpu-deficient HIV-1 (HIV_LAI_Δvpu) in IFN-treated Tetherin-KO cells shows increased virus release as compared to control cells (Tetherin_1 = 20.5-fold, Tetherin_2 = 14-fold in Figure 4E - left panel), confirming the late inhibition of Vpu-deficient HIV by Tetherin. Infection of these cell pools with wt HIV also shows significantly-increased virus release, suggesting that HIV-1_LAI_ Vpu does not completely antagonize IFN-induced Tetherin in THP-1 cells (Tetherin_1 = 4.4-fold, Tetherin_2 = 2.14-fold in Figure 4E – right panel). Therefore, Tetherin is a late-acting ISG contributing to IFN inhibition of HIV-1_LAI_ in THP-1 cells.

### The HIV-CRISPR screen also identifies HIV dependency factors

Although we designed our screen specifically to find IFN-induced factors restricting HIV-1 in THP-1 cells, HIV-CRISPR screening can also identify HIV dependency factors. The sgRNA sequences of genes that HIV uses for enhanced viral replication will be depleted in viral supernatants as the virus will be less well able to replicate specifically in these cells (Figure 1C). Analysis of the negative MAGeCK Gene Scores, representing genes for which sgRNAs were depleted in HIV supernatants, identifies a panel of candidate host factors targeted by the PIKA_HIV_ library that are important for HIV replication (Figure 5A). The top hit is the HIV-1 co-receptor CXCR4 (Figure 5A) which is required for entry by HIV-1_LAI_ (note: sgRNAs targeting the receptor, CD4, are not present in the PIKA_HIV_ library). The next highest scoring hit is Siglec-1/CD169, an HIV attachment factor that has been characterized to facilitate *trans* infection of CD4+ T cells by DCs through binding to sialylated glycosphingolipids on the HIV particle (Izquierdo-Useros et al., 2012; Puryear et al., 2013) (Figure 5A). CD169 is upregulated by IFNα in THP-1 cells (Figure 5B, left panel). Our screen only assays cell-autonomous effects suggesting that CD169 also plays a role in *cis*-infection of monocytic cells, consistent with recent work showing enhanced infection of THP-1 cells by CD169, specifically in the presence of IFNα (Akiyama et al., 2017). Indeed, when CD169 expression is knocked-down (Figure 5B middle panel) these cells are less susceptible to infection both in the presence and absence of IFN pretreatment (Figure 5B right panel), although this effect is stronger in presence of IFNα (6.5-fold vs 4.7-fold; Figure 5B right panel). Thus, we find that Siglec-1/CD169 is an IFN-induced, HIV dependency factor in THP-1 cells.

**FIGURE 5.**
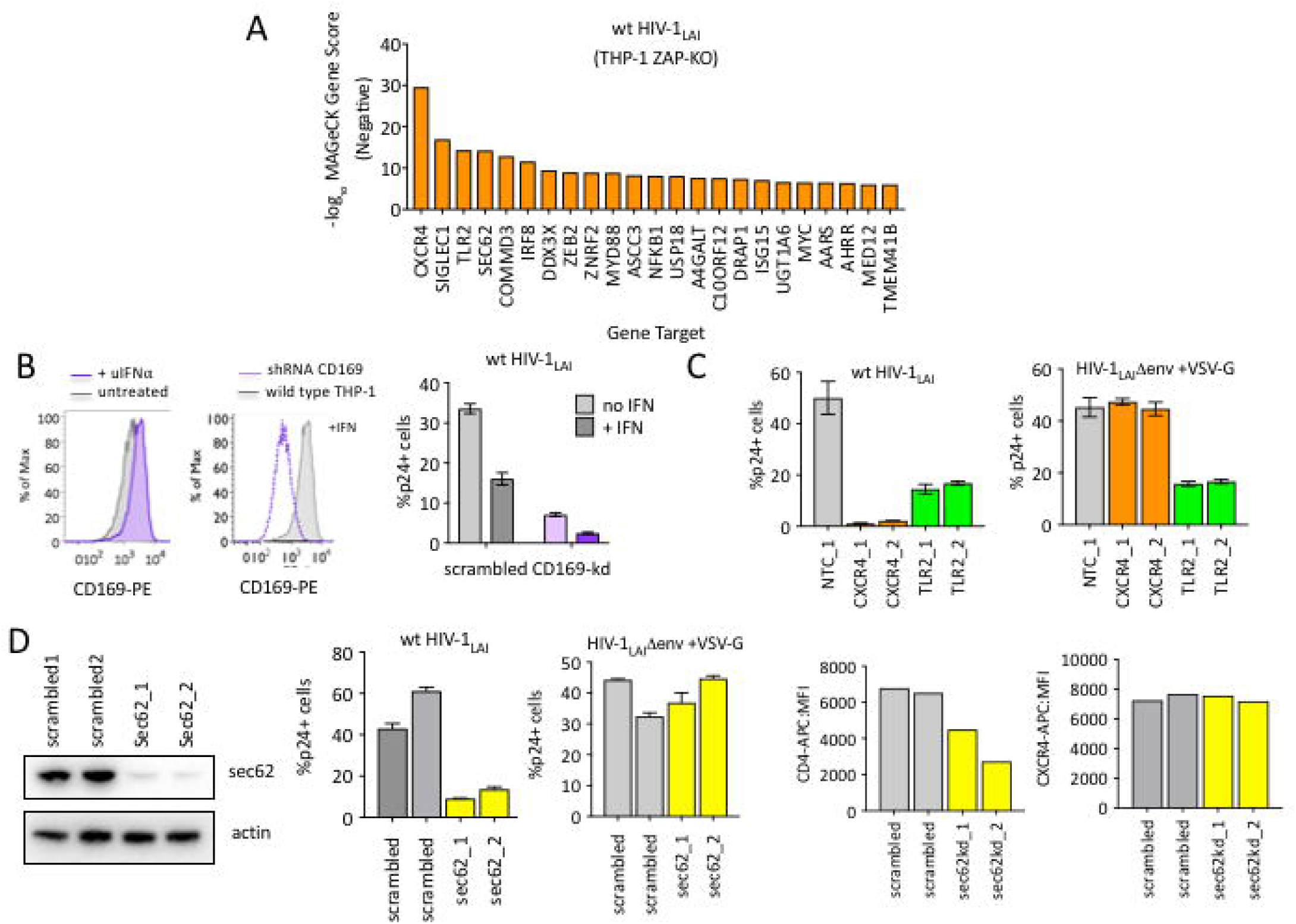
HIV-CRISPR Screening Identifies HIV Dependency Factors. <u>A</u>: Negative MAGeCK Gene Scores across both ZAP-KO Screens ranked from most depleted genes on the X-axis. Only the top 25 hits are shown. <u>B</u>: <u>Left:</u> THP-1 cells were stimulated overnight with IFNα and assayed for cell surface SIGLEC1/CD169 expression by flow cytometry. <u>Middle</u>: Control (scrambled - gray) THP-1 cells andTHP-1 cells transduced with a SIGLEC1/CD169-targeting shRNA construct (dotted purple line) were assayed for cell surface SIGLEC1/CD169 expression after overnight IFNα treatment. <u>Right</u>: Infection of control (gray - wild type) and SIGLEC1/CD169 knockdown THP-1 (purple - CD169-KD) with and without IFNα (1000 U/mL u IFNα) and assayed by intracellular p24gag 2 days after infection <u>C</u>: Infection of control (gray - NTC), CXCR4-KO pools (orange) and TLR2-KO pools (green) were assayed for the % of cells expressing HIV p24gag 2 days post-infection by intracellular staining and flow cytometry. <u>Left</u>: wt HIV-1_LAI_ (n=3). <u>Right</u>: HIV-1_LAI_Denv+ VSV-G (n=3). <u>D</u>: <u>Left</u>: SEC62 knockdown after transduction with two LKO SEC62 shRNA constructs. Western blot of the sec62-targeting shRNA cell lines is shown together with 2 control (scrambled) cell lines. Loading control = actin. <u>Left/Middle and Middle</u>: Infection of SEC62-KD (orange) and control (scrambled-KD cells (gray) with wt HIV-1_LAI_ or HIV-1_LAI_Δenv + VSV-G. The % of cells expressing HIV p24gag 2 days post-infection is shown. <u>Right-middle and Right</u>: The mean florescence intensity (MFI) of CD4 and CXCR4 cell surface staining of control (scrambled) and SEC62-KD THP-1 cell pools.

TLR2, a toll-like receptor characterized to recognize bacterial PAMPs (Akira, Uematsu, & Takeuchi, 2006) is the next highest-scoring hit in our dependency factor analysis. We infected TLR2-KO cell pools in tandem with NTC (negative control) and CXCR4-KO (positive control) THP-1 cell pools generated by transduction with lentiCRISPRv2 sgRNA constructs. Infection of these cells demonstrates lower infection as compared to the controls, although this effect is not as extreme as for CXCR4 (Figure 5C left panel; 31-fold decreased infection for CXCR4, 3-fold for TLR2). Of note, infection with VSV-G pseudotyped HIV-1 shows a loss of infectivity similar to wild-type HIV-1, suggesting that the effect of TLR2 on enhanced infection is independent of viral entry (Figure 5C right panel). Finally, we investigated the effect of SEC62, on HIV replication. SEC62 is a component of the protein translocation machinery in the ER membrane. Knockdown of SEC62 by transducing THP-1 cells with two lentiviral shRNA constructs targeting SEC62 shows significant loss of expression as measured by Western Blot (Figure 5D). Infection of these cells with wt HIV-1 shows decreased levels of infection (Figure 5D), Therefore, SEC62 is a dependency factor for HIV replication in THP-1 cells. As SEC62 is a component of the machinery that mediates translocation of transmembrane proteins into the ER membrane for targeting to the cell surface, we reasoned that SEC62 knockdown may be affecting cell-surface expression of HIV receptors, co-receptors or other cell-surface markers mediating attachment and/or entry of HIV. Consistent with this hypothesis, infection via an alternative entry pathway via pseudotyping HIV-1 particles with VSV-G, demonstrates equivalent infection in control and SEC62 knockdown cells (Figure 5D). Analysis of the cell surface expression of the HIV-1 receptor, CD4, shows that levels of CD4 on the cell surface are decreased in SEC62 knockdown cells (Figure 5D). Interestingly, we do not observe decreased cell-surface expression of CXCR4 (Figure 5D), suggesting that the effect of SEC62 knockdown on cell surface proteins is not global but specific to certain transmembrane proteins. In summary, HIV-CRISPR screening can identify HIV host dependency factors in addition to restriction factors.

## Discussion

We have designed and validated a novel CRISPR knockout screening approach in which the virus itself serves to report levels of infection to identify genes important for HIV infection. Subsequent deep sequencing can quantitate effects of individual gene knockouts on viral replication in a massively-parallel fashion. Using the PIKA_HIV_ library targeting human ISGs, we demonstrated that the IFN-mediated inhibition of HIV in THP-1 cells is due to the combined action of a small panel of ISGs that includes known HIV restriction factors like MxB, TRIM5alpha, IFITM1 and Tetherin. Each of these ISGs individually imposes only modest restriction of HIV replication but together mediate robust restriction of HIV replication. While HIV has evolved antagonism or evasion strategies for restriction factors that limit replication in cells from other species, the results here imply that even well-adapted HIV strains, such as HIV_LAI_, are not able to completely antagonize or escape some host encoded restriction factors. Such incomplete antagonism may be due to conflicting evolutionary pressures acting on the HIV genome.

### The HIV-CRISPR Screening Approach

Other high-throughput screens with siRNA pools have focused on identifying dependency factors that the virus takes advantage of to infect cells (Brass et al., 2008; Konig et al., 2008; Zhou et al., 2008). Similarly, a recent CRISPR knockout screen demonstrated that a pooled CRISPR approach to gene knockout could identify HIV dependency factors (Park et al., 2017). Importantly, all of these high-throughput approaches, with the exception of the recent CRISPR knockout screen, still require individual gene knockdowns or overexpression in individual wells. The HIV-CRISPR screening approach represents a significant advance in screening for host factors that affect HIV replication in several ways, including: (1) we can simultaneously screen thousands of gene targets in a single experiment, (2) we can use any virus strain, (3) we do not need any type of reporter to assay infections as virus replication itself provides the assay readout and (4) we can capture host factors that affect all stages of the HIV life-cycle including entry, nuclear import, integration, transcription, nuclear export, translation, packaging, budding and release. After finding ZAP-mediated inhibition of the HIV-CRISPR vector used in our screening approach, we modified our PIKA_HIV_ screen to avoid this inhibition by specific KO of ZAP expression and rescreening in ZAP-KO THP-1 clonal lines. Genetic deletion of ZAP resulted in enhanced performance of the HIV-CRISPR screen and allowed for our identification of ISGs contributing to the IFN block in THP-1 cells. Further the data presented here demonstrates that the screen is sensitive enough to find key factors in just a single round of viral replication, even when multiple factors together mediate potent inhibition.

### Incomplete antagonism of HIV restriction factors

Our finding of significant TRIM5alpha restriction in human cells suggests that HIV is still partially-sensitive to TRIM5alpha-mediated restriction. Similarly, we find that IFN-induced MxB restricts infection in THP-1 cells, consistent with previous work (Goujon et al., 2013; Kane et al., 2013; Z. Liu et al., 2013). Both TRIM5 and Tetherin are rapidly-evolving genes in primates with described consequences for host adaptation by primate lentiviruses (Lim, Malik, & Emerman, 2010; H. L. Liu et al., 2005). Capsids from diverse primate lentiviruses have adapted to TRIM5 alleles in various primates and are variably-restricted by TRIM5alpha orthologs (Kirmaier et al., 2010). Selection for capsid mutations that evade TRIM5alpha restriction is a key adaptive step that HIV and related SIVs must make to successfully replicate in a particular primate species (F. Wu et al., 2013). Consistent with a role for TRIM5alpha in humans, TRIM5alpha is active against HIV in Langerhans cells (Ribeiro et al., 2016). Further, CA mutations in HIV-infected individuals have been associated with sensitivity to TRIM5alpha restriction (Battivelli et al., 2010; Onyango et al., 2010). Also of note, most studies of HIV have not been done in the context of IFN despite evidence that TRIM5alpha is highly-IFN upregulated in HIV target cells (Carthagena et al., 2009). Thus, our finding of IFN-mediated TRIM5alpha inhibition of HIV represents a potentially-important role of TRIM5alpha particularly during acute infection when IFN levels are high. More stable artificial variants of human TRIM5alpha can inhibit HIV-1 (Richardson, Guo, Xin, Yang, & Riley, 2014), suggesting that increased TRIM5 levels, such as after IFN induction, may play a role in restricting HIV replication. Evasion of TRIM5alpha restriction may come at the cost of loss of fitness due to other requirements for capsid function within host cells. Key capsid-mediated functions include uncoating, nuclear import and integration. Further, capsid sequences also mediate evasion of other restriction factors, including MxB, or escape from host CTL responses. Primary isolates of HIV-1 have increased sensitivity to TRIM5alpha that is proposed to be driven by CTL escape variants (Battivelli et al., 2011). Therefore, we speculate that this TRIM5alpha sensitivity may underscore the requirement of HIV proteins to balance multiple functions simultaneously to infect human cells.

Similar to TRIM5, our finding of inhibition of wt HIV by Tetherin despite intact Vpu expression may suggest a functional tradeoff in which HIV Vpu is unable to completely antagonize host cell Tetherin activity. Adaptation HIV-1 group M to the unique form of human Tetherin allele required evolution of the viral protein Vpu to antagonize Tetherin (Lim et al., 2010; Sauter et al., 2009). Consistent with conflicting evolutionary constraints, IFN treatment in HCV-and HIV-coinfected patients resulted in evolution of Vpu variants with stronger Tetherin antagonism when ISGs are expressed *in vivo* (Pillai et al., 2012). Perhaps more complete antagonism of Tetherin by Vpu would compromise some of the other functions of Vpu in cells (Apps et al., 2016; Margottin et al., 1998; Schubert et al., 1998; Shah et al., 2010). Further, a moderate level of Tetherin antagonism could be selected for if cell-to-cell transmission is enhanced by Tetherin restriction (Gummuluru, Kinsey, & Emerman, 2000; Jolly, Booth, & Neil, 2010), such as is observed for MoMLV in mice (Liberatore, Mastrocola, Powell, & Bieniasz, 2017). Like TRIM5 and Tetherin, escape from IFITMs also appears to be subject to conflicting evolutionary pressures. IFITMs may exert significant selective pressure *in vivo* as HIV evolves increased susceptibility to IFITMs over the course of infection (Foster et al., 2016). A similar example of incomplete antagonism of human restriction factors can be found in HIV-infected patients in which a signature of APOBEC3 G-to-A hypermutation in integrated proviruses can be observed (Cuevas, Geller, Garijo, Lopez-Aldeguer, & Sanjuan, 2015; Sadler, Stenglein, Harris, & Mansky, 2010) despite the fact that HIV encodes an antagonist, Vif, that targets APOBEC3 proteins for degradation. Finally, it may be that further adaptation of HIV for efficient replication in human cells may not be possible due to constraints on viral evolution imposed by concurrently-acting restriction factor barriers.

### Additional novel antiviral phenotypes

We found that ZAP mediates a small, but detectable inhibition of HIV replication as we find enhanced infection of ZAP-KO cells both in the presence and absence of IFN pretreatment (Supplemental Figure S2). Similar to the effect of ZAP, N4BP1 (Nedd4-binding protein 1) also has a modest effect on HIV replication both after IFN pretreatment and when constitutively-expressed (Supplemental Figure S2). In our screen, the anti-lentiviral function of N4BP1 appears to be genetically linked to ZAP activity as N4BP1 is no longer a hit in the ZAP-KO screen (Supplemental Table S4). Therefore, N4BP1 may modify or enhance ZAP-mediated antiviral activity similar to the modification of ZAP activity described for TRIM25 (M. M. Li et al., 2017). In addition to ZAP, TRIM25 and N4BP1, several other ISGs scored highly in the PIKA_HIV_ screen including UBE2L6 and LGALS3BP (also known as 90K or M2BP). UBE2L6 and 90K inhibit HIV in over-expression assays (Jain et al., 2018; Lodermeyer et al., 2013; Q. Wang, Zhang, Han, Wang, & Gao, 2016) and SAMD9L was recently shown to be an IFN-induced restriction factor for poxviruses (Meng et al., 2018).

### HIV-CRISPR screening can identify HIV dependency factors

Despite targeting less than 10% of the genes in the human genome by our PIKA_HIV_ library, we were able to identify and validate a small panel of HIV dependency factors that HIV usurps to enhance infection in THP-1 cells. We demonstrate that the HIV attachment factor, SIGLEC1/CD169, plays a role in enhancing infection in THP-1 cells *in cis* rather than the more fully described role of SIGLEC1 to mediate infection from dendritic cells to T cells *in trans* (Izquierdo-Useros et al., 2012; Puryear et al., 2013). Further, we find that TLR2 mediates enhanced infection of THP-1 cells by HIV-1 regardless of viral entry pathway used, as it impacted infection through both the HIV envelope and the VSV-G glycoprotein (Figure 5C). Recent work in CD4+ T cells has similarly demonstrated enhanced infection and/or viral production in T cells on stimulation of TLR2 (Bolduc, Ouellet, Hany, & Tremblay, 2017; Ding & Chang, 2012; Ding et al., 2010; Equils et al., 2003; Henrick, Yao, Rosenthal, & team, 2015). Of note, MYD88, a downstream effector for TLR2 activation of transcription, is also a strong hit in our dependency factor screening (Figure 5A) suggesting that it is the downstream signaling functions of TLR2 that are important for enhancing infection. In addition to CD169 and TLR2, our identification of SEC62 as a novel HIV dependency factor that correlates with CD4 receptor cell surface expression highlights the ability of the HIV-CRISPR screening approach to find genes that function in pathways (such as CD4 receptor expression) important for HIV infection. A recently-published HIV CRISPR screen (Park et al., 2017) to uncover important HIV host factors differs from our study in using Tat-driven LTR-GFP reporter gene expression as well as many rounds of spreading infection across multiple weeks in culture. In contrast, PIKA_HIV_ screening is performed over a single round of infection in three days and the screen relies on virus replication itself to enrich for gene targets of interest. Consistent with these key differences in approach, the dependency factor genes we identify in the PIKA_HIV_ screen differ from the findings of Park *et. al*. Both screens do identify the appropriate HIV co-receptor (CXCR4 in our study and CCR5 in the Park *et al*. study). Of note, 3 of the 5 genes identified by Park *et al*. (TPST2, SLC35B2 and CD4) are not represented in the PIKA_HIV_ library and, therefore, could not be identified in our screen. Further studies using whole genome CRISPR libraries for the HIV-CRISPR approach should identify further HIV dependency factors that were not present in our current PIKA_HIV_ library.

In summary, we developed a novel screen that is highly sensitive to detect restriction factors for HIV-1. This new tool shows that the IFN inhibition of HIV-1 in a monocytic cell line is due the combined function of fewer than 8 different genes. Our results demonstrate that IFN-mediated inhibition of HIV-1 in THP-1 cells is mediated by restriction factors for which HIV has described mechanisms of antagonism and/or escape. The increased IFN sensitivity of specific HIV strains, such as those isolated during chronic HIV infection, may due to relaxation of constraints on the virus that would otherwise limit virus replication during transmission events. We propose that conflicting functional constraints acting on HIV may result in incomplete antagonism or escape from host ISGs during chronic infection.

## Acknowledgments

We thank Patrick Paddison, Phil Corrin, Lucas Carter, Yu Ding, Pia Hoellerbauer, Dan Kuppers for assembly of the PIKA library and assistance with screening methodology, Stephanie Rainwater and Abby Felton for technical assistance, Chris Large for a parallel screen using the PIKA library for genes necessary for induction of ISGs and the Fred Hutch Shared Resources Bioinformatics and Genomics Cores (NCI 5 P30 CA015704-43) and Harmit Malik for discussions and comments on the manuscript. This work was supported by NIH grant R01 AI30927 (M.E), a CCEH Pilot Grant P30 DK56465 (M.O.), UW/FHCRC CFAR New Investigator Award P30 AI027757 (M.O.) and a Belgian American Educational Foundation Fellowship (J.V.). Construction of the PIKA library was supported by DP1 DA039543 to Julie Overbaugh.

The authors have no competing interests.

## Materials and Methods

### Interferon-Stimulated Gene Dataset

1905 human ISGs were selected from gene expression datasets of type I IFN-stimulated cells (Goujon et al., 2013; Hung et al., 2015; Linsley, Speake, Whalen, & Chaussabel, 2014) or from previously assembled ISG overexpression (Schoggins et al., 2011) or shRNA libraries (J. Li et al., 2013). These included all the genes from the previously assembled ISG libraries (J. Li et al., 2013; Schoggins et al., 2011) as well as additional ISGs as defined here. For the GSE46599 dataset (Goujon et al., 2013), raw probe-level signal intensities from Illumina HumanHT-12 V4.0 expression BeadChip data were retrieved from GEO, then background-corrected, quantile-normalized and log_2_-transformed using the Bioconductor package lumi (Du, Kibbe, & Lin, 2008). Fold changes (FC) in expression between type I IFN-treated and untreated samples were calculated for untreated and PMA-treated THP-1 cells, primary CD4+ T cells and primary macrophages. For THP-1 cells, genes with FC ≥ 2 were selected. For primary cells, genes with a donor-specific FC ≥ 2 in at least 2 out of 3 donors were selected. For the GSE60424 dataset (Linsley et al., 2014), TMM normalized RNA-seq read count data (Illumina HiScan) were retrieved from GEO. FC in expression in whole blood, isolated CD4+ T cells and monocytes of a Multiple Sclerosis patient, pre- and post-treatment with AVONEX (IFNβ), were calculated and genes with FC ≥ 2 were selected. For the GSE72502 dataset (Hung et al., 2015), *de novo* identification of differentially-expressed genes in IFNα treated PBMCs was performed from the raw RNA sequencing data (Illumina Genome Analyzer). SRA files were retrieved from GEO and converted to FASTQ format using NCBI’s SRA toolkit. Reads were mapped to the human reference genome (hg19) using GSNAP (T. D. Wu, Reeder, Lawrence, Becker, & Brauer, 2016) and quantified using HTSeq (Anders, Pyl, & Huber, 2015). Differentially-expressed (DE) genes were identified using the Bioconductor edgeR package (Robinson, McCarthy, & Smyth, 2010). DE genes were defined at an FDR threshold of 0.05. The glmTreat function was used to detect genes with a FC significantly greater than 1 between the IFN-treated and control samples. Finally, non-coding RNAs and pseudogenes were removed from the list. Inspection of the curated list of genes showed that overlap between the different datasets was limited and many genes (> 2000) were only present in 1 of the 10 datasets/libraries. As such, a second selection round was performed in which the expression threshold for genes present in only one of the datasets was raised to FC≥3. For genes present in at least two datasets, the initial cut-off of FC≥2 was kept. Finally, 35 additional genes identified through RNA sequencing gene expression analysis as being responsive to both type I/type III IFN and IL-1β were also included (M. Gale, personal communication, October, 2015). For analysis of IFN induction specific to THP-1 cells, raw signal intensities were downloaded from GEO (GSE46599) and the data was quantile normalized using the Bioconductor package lumi. For a given probe, both samples from at one least condition were required to have a detection p-value <= 0.05. The Bioconductor package limma was used to identify significantly differentially expressed probes. A false discovery rate (FDR) method was employed to correct for multiple testing (Reiner, Yekutieli, & Benjamini, 2003), with differential expression defined as |log2 (ratio)| ≥ 0.585 (± 1.5-fold) with the FDR set to 5%.

### Cell Culture

The THP-1 monocytic cell line (ATCC) was cultured in RPMI (Invitrogen) with 10% FBS, Pen/Strep, 10mM HEPES, 0.11 g/L sodium pyruvate, 4.5 g/L D-Glucose and Glutamax. 293T and TZM-bl cells were cultured in DMEM (Invitrogen) with 10% FBS and Pen/Strep. For some validation studies, THP-1 cells with single-cell sorted into 96-well plates to create individual clonal lines (BD FACS Aria II - Fred Hutch Flow Cytometry Core). Universal Type I Interferon Alpha was obtained from PBL Assay Science (Catalog No. 11200-2), diluted to 10^5^ Units/mL in sterile-filtered PBS/1% BSA according to the activity reported by manufacturer and frozen in aliquots at −80°C. All Puromycin selections were done at 0.5-1ug/mL.

### Plasmids

lentiCRISPRv2 plasmid was a gift from Feng Zhang (Addgene #52961). lentiCRISPRv2-mCherry was a gift from Agata Smogorzewska (Addgene # 99154). pMD2.G and psPAX2 were gifts from Didier Trono (Addgene #12259/12260). lentiCRISPRv2 constructs targeting genes of interest were cloned into BsmBI-digested lentiCRISPRv2 by annealing complementary oligos (Supplemental Table S5) with overhangs that allow directional cloning into lentiCRISPRv2. Stable LKO SEC62 shRNA lentiviral vectors were obtained from Sigma. SEC62_1:

CCGGCCAGGAAATCATGGAACAGAACTCGAGTTCTGTTCCAT GATTTCCTGGTTTTTG

(TRCN0000289739). SEC62_2:

CCGGGAAATGAGAGTAGGTGTTTATCTCGAGATAAACACCTACTCT CATTTCTTTTTG

(TRCN0000289833). Scramble shRNA

(CCTAAGGTTAAGTCGCCCTCGCTCGAGCGAGGGCGACTTAACCTTAGG) was a gift from David Sabatini (Addgene #1864). The CD169 shRNA (Sigma TRCN155147)

(CCGGGTGTGGAGATTCACAACCCTTCTCGAGAAGGGTTGTGAATCTCCACACTTTTTTG) was a gift from Rahm Gummuluru. The CD169 shRNA sequence was subcloned into pLKO.1neo (Addgene #13425) using EcoRI and AgeI sites. HIV-CRISPR was constructed (Genscript) by inserting a synthesized 433bp sequence from HIV-1_LAI_ into the deleted 3’ LTR U3 sequence of lentiCRISPRv2. HIV-1_LAI_ LT insert:

ATCCTTGATCTGTGGATCTACCACACACAAGGCTACTTCCCTGATTGGCAGAACTACACACCAGGGCCAGGGGTCAGATATCCACTGACCTTTGGATGGTGCTACAAGCTAGTACCAGTTGAGCCAGATAAGGTAGAAGAGGCCAATAAAGGAGAGAACACCAGCTTGTTACACCCTGTGAGCCTGCATGGAATGGATGACCCTGAGAGAGAAGTGTTAGAGTGGAGGTTTGACAGCCGCCTAGCATTTCATCACGTGGCCCGAG AGCTGCATCCGGAGTACTTCAAGAACTGCTGACATCGAGCTTGCTACAAGGGACTTTCCGCTGGGGACTTTCCAGGGAGGCGTGGCCTGGGCGGGACTGGGGAGTGGCGAGCCCTCAGATGCTGCATATAAGCAGCTGCTTTTTGCCTGTACTGGGTCTCTCTGGTTA. The wild type (HIV-1_LAI_), env-deleted (HIV-1_LAI_ VSV-G) and *vpu*-deficient (HIV_LAI_Δvpu = VpuFS/Rap5) HIV-1_LAI_ proviruses were previously described (Bartz & Vodicka, 1997; Gummuluru et al., 2000; Peden, Emerman, & Montagnier, 1991).

### ISG CRISPR/Cas9 sgRNA Library Construction

4 sgRNA sequences were selected randomly from the Brunello library for each gene target (Doench et al., 2016) and 4 additional non-identical sgRNAs were subsequently selected randomly from the Genome-scale CRISPR Knock-Out (GeCKO v2) library (Sanjana, Shalem, & Zhang, 2014a; Shalem, Sanjana, Hartenian, Shi, Scott, Mikkelsen, et al., 2014). For genes for which 8 unique sgRNAs could not be obtained from these libraries, additional sgRNAs were added from the Moffat (Hart et al., 2015) and Sabatini/Lander libraries (T. Wang et al., 2015; T. Wang et al., 2014). 12 genes contained no sgRNAs in any of the libraries and for those genes 8 new sgRNAs were designed using the sgRNA *Designer* from the *Broad Institute* (http://portals.broadinstitute.org/gpp/public/analysis-tools/sgrna-design). A total of 15,348 unique sgRNA sequences were synthesized. The sgRNAs were split in two pools for synthesis (4 per gene in each pool) and two independent sets of 200 Non-Targeting Control (NTC) sgRNAs obtained from the GeCKOv2 library were added in duplicate to each pool. The PIKA_HIV_ ISG-targeting sgRNA library was synthesized (Twist Biosciences) and cloned into HIV-CRISPR. Oligo pools were amplified using Phusion HF (Thermo) using 1 ng of pooled oligo template, primers ArrayF and ArrayR (ArrayF primer:TAACTTGAAAGTATTTCGATTTCTTGGCTTTATATATCTTGTGGAAAGGACGAAACACCG and ArrayR primer:ACTTTTTCAAGTTGATAACGGACTAGCCTTATTTTAACTTGCTATTTCTAGCTCTAAAAC), an annealing temperature of 59°C, an extension time of 20s, and 25 cycles. Following PCR amplification, a 140bp amplicon was gel-purified and cloned into BsmBI digested vectors using Gibson assembly (NEB). Each Gibson reaction was carried out at 50°C for 60 minutes in a thermocycler. 1μl of the reaction was used to transform 25μl of electrocompetent cells (Stellar Competent Cells; Clontech) according to the manufacturer’s protocol using a GenePulser (BioRad). To ensure adequate representation, sufficient parallel transformations were performed and plated onto ampicillin containing LB agarose 245 mm × 245 mm plates (Thermo Fisher) at 200-times the total number of oligos of each library pool. After overnight growth at 37°C, colonies were scraped off, pelleted, and used for plasmid DNA preps using the Endotoxin-Free Nucleobond Plasmid Midiprep kit (Takara Bio #740422.10).

### Virus and lentivirus production

293T cells (ATCC) were plated at 2×10^5^ cells/mL in 2mL in 6-well plates one day prior to transfection using TransIT-LT1 reagent (Mirus Bio LLC) with 3μL of transfection reagent per μg of DNA. For lentiviral preps, 293Ts were transfected with 667ng lentiviral plasmid, 500ng psPAX2 and 333ng MD2G. For HIV-1 production, 293Ts were transfected with 1ug/well proviral DNA. One day post-transfection media was replaced. Two or three days post-transfection viral supernatants were clarified by centrifugation (1000g) and filtered through a 20μm filter. For PIKA_HIV_ library preps, supernatants from 40 x 6-well plates were combined and concentrated by ultracentrifugation. 30mL of supernatant per SW-28 tube were underlaid with sterile-filtered 20% sucrose (1mM EDTA, 20mM HEPES, 100mM NaCl, 20% sucrose) and spun in an SW28 rotor at 23,000rpm for 1 hour at 4°C in a Beckman Coulter Optima L-90K Ultracentrifuge. Supernatants were decanted, pellets resuspended in DMEM over several hours at 4°C and aliquots frozen at −80°C. All viral and lentiviral infections and transductions were done in the presence of 20μg/mL DEAE-Dextran (Sigma; D9885).

### PIKA_HIV_ Screening

Large-scale preps of the PIKA_HIV_ lentiviral library were titered by a colony-forming assay in TZMbl cells and used to transduce THP-1 cells at an MOI of 0.7. Cells were selected in Puromycin (0.5 μg/mL) for 10 - 14 days. 8×10^6^ cells per replicate (>500X coverage of the PIKA_HIV_ library) were infected at a viral dose determined to allow approximately 50% of cells in culture to be infected by spinoculation at 1100×_g_ for 30 minutes with 20μg/mL DEAE-Dextran. After overnight incubation, cells were resuspended in media with or without IFNα at 5×10^5^ cells/mL. Cells and supernatants were collected 3 days post infection. Genomic DNA was extracted from cell pellets with a QIAamp DNA Blood Midi Kit (Qiagen 51183) and genomic DNA eluted in water. Viral supernatants were spun at 1100×_g_ to remove cell debris, filtered through a 0.2μm filter, overlaid on a 20% sucrose cushion and concentrated in SW28 rotor for 1 hour at 4°C. After resuspension in PBS, viral RNA was extracted (QIAamp viral RNA Kit, Qiagen, 52904). sgRNA sequences present in the genomic DNA and viral supernatants were amplified by PCR and RT-PCR, respectively, using primers specific for the HIV-CRISPR construct (Supplemental Table S5) (Toledo et al., 2015). Libraries were then barcoded/prepared for Illumina sequencing by a second round of PCR (Supplemental Table S5). Each amplicon was then cleaned up through double-sided SPRI (Agencourt AMPure XP Beads – Beckman Coulter #A63880), quantitated with a Qubit dsDNA HS Assay Kit (Q32854 – ThermoFisher) and pooled to 2nm for each library. Pooled, multiplexed libraries were then sequenced on a single lane of an Illumina HiSeq 2500 in Rapid Run mode (Fred Hutch Genomics and Bioinformatics Shared Resource).

### Screen analysis

Following demultiplexing of libraries to assign sequences to each sample (allowing no mismatches), reads were trimmed and aligned to the PIKA_HIV_ sgRNA library, using Bowtie (Langmead, Trapnell, Pop, & Salzberg, 2009). NTC sgRNA sequences were iteratively binned to create an NTC sgRNA set as large as the ISG gene set in the PIKA_HIV_ library. Relative enrichment of sgRNAs and genes were analyzed using the MAGeCK statistical package (W. Li et al., 2014). For the VSV-G screen a single IFITM1-targeting sgRNA sequence (AGCATTCGCCTACTCCGTGA) with complete homology to IFITM3 was removed from the analysis.

### Sliding window analysis of CG dinucleotide content

An Excel (Microsoft) worksheet was created to analyze the CG dinucleotide content of the HIV-CRISPR-NTC1 transcript from the beginning of the 5’R region to the end of the 3’R region. The HIV-CRISPR sequence was broken into fragments of 3 nucleotides (codons), and at each position the number of CG dinucleotides within or between two adjacent codons was determined. The CG counts at each position over the length of the sequence were then summed within a sliding window of 67 codons (201 nucleotides) and plotted against the nucleotide position of the transcript in GraphPad Prism.

### Digital droplet PCR (ddPCR)

Wild-type or ZAP-knockout THP-1 cells were transduced with a pooled library of HIV-CRISPR encoding 39 distinct gRNAs at an MOI of 0.5 and selected with puromycin for 15 days as described above. Cells were infected with HIV-1_LAI_ at an MOI of 1. Three days post-infection, viral supernatants were cleared by centrifugation, filtered through a 0.4μm filter, and viral RNA was extracted from 140μL of supernatant using the QIAamp viral RNA Kit, with subsequent aliquoting and freezing at −80°C. cDNA was synthesized from viral RNA with random hexamers, and the number of copies of either HIV or HIV-CRISPR per μL of cDNA was quantified by ddPCR using the QX200™ Droplet Digital™ PCR System (Bio-Rad, Hercules, CA). HIV was detected using previously published primers and probe directed towards *pol* (Benki et al., 2006). To specifically detect HIV-CRISPR, we used primers ddPCR-cPPT-F (GTA CAG TGC AGG GGA AAG), ddPCR-U6-R (ATG GGA AAT AGG CCC TCG), and probe cPPT-probe (6-FAM/ZEN-AGA CAT AAT AGC AAC AGA CAT ACA AAC-IBFQ) (Integrated DNA Technologies, Skokie, IL). Both sets of reactions were set up according to the manufacturers protocols with an annealing temperature of 60°C. The cPPT-U6 primers were found to be specific to HIV-CRISPR, as no amplification was detected in untransduced cells infected with HIV-1. Control reactions on viral RNA without reverse transcriptase revealed that carry-over plasmid contamination from viral preps accounted for only a low level (<50 copies/μL) of amplification.

### Knockout and Knockdown Cell Pools and Clones

ZAP knockout cell pools were created by electroporating THP-1 cells with a custom ZAP-targeting crRNA (ATGTGGAGTCTTGAACACGG; IDT). 1μL crRNA was resuspended at 160μM in 10mM Tris pH 7.4 and complexed at an equimolar ratio with 1μL 160μM tracrRNA (IDT #1072534) and incubated 30 minutes at 37°C followed by addition of 2μL of 40μM Cas9-NLS (UC Berkeley MacroLab) and further incubation at 37°C for 15 minutes to create the ZAP-targeting crRNP complexes. 3.5μL crRNP was added to 5×10^5^ THP-1 cells resuspended in Amaxa SG Cell Line 96-well Nucleofector Kit (Lonza #V4SC-3096) and electroporated according to the manufacturer’s protocol (Lonza 4D Nucleofector). 80μL of prewarmed media was added, followed by incubation for 30-minute recovery in the 37°C incubator. Cells were then resuspended at 2.5×10^5^ cells/mL in 500μL in a 24-well plate for 48 hours before single cell sorting into 96 well U-bottom plates containing RPMI media supplemented with 20% FBS (BD FACS Aria II - Fred Hutch Flow Cytometry Core). MxB-KO clonal lines were generated by transduction with lentiCRISPRv2 containing MxB-targeting sgRNA sequences (see Supplemental Table S5 for sgRNA sequences) followed by single-cell cloning and puromycin selection. lentiCRISPRv2 KO cell pools targeting STAT1, STAT2, IRF9, MxB, TRIM5, IFITM1, Tetherin, CXCR4 and TLR2 as well as two Non-Targeting Controls (NTC_1 and NTC_2) were created through transduction with lentivirus and selection in Puromycin (see Supplemental Table S5 for sgRNA sequences). Both KO cell pools and individual KO cell lines were validated using Western blotting, flow cytometry and/or genomic editing analysis as described below. shRNA knockdown cell pools were made by transducing wildtype THP-1 cells with lentiCRISPRv2 shRNA constructs and selected in RPMI containing 1μg/mL Puromycin for two weeks prior to validating via Western blotting or flow cytometry.

### Genomic Editing Analysis

Knockout cells were harvested and either lysed in Epicentre QuickExtract DNA Extraction Solution (Lucigen QE09050) for direct PCR amplification or genomic DNA was extracted (QIAamp DNA Blood Mini Kit – Qiagen #51185). Edited loci were amplified from cell pool DNA using primers specific to each targeted locus as previously published (Hultquist et al., 2016). PCR amplicons were sequenced (Fred Hutch Shared Resources Genomics Core – sanger sequencing) and analyzed by ICE (Synthego) to determine the percent of alleles edited at each locus in the cell population (Hsiau et al., 2018). Editing was confirmed at each locus (Supplementary Table S6; KO scores varied across pools from 48% to 93%).

### Antibodies

For Western blotting the following antibodies were used as follows: MxB (Santa Cruz sc-271527) at 1:200, Sec62 (Abcam ab168843) at 1:2000 and actin (Sigma A2066) at 1:5000. Secondary antibodies were used as follows: 1:5000 donkey anti-goat IgG-HRP (Santa Cruz Biotechnology sc-2020) and 1:5000 goat anti-rabbit IgG-HRP (Santa Cruz Biotechnology sc-2004). For flow cytometry, antibodies were used as follows: CD4 (BD Pharmingen 555349) 1:50, CXCR4 (eBioscience 17-9999-42) 1:50, CD-169 (BioLegend 346003) 1:50, TLR2 (BioLegend 309707) 1:100, Tetherin (BioLegend 348410) 1:50.

### Flow Cytometry

For intracellular Gag_p24_ (p24) staining, cells were harvested and fixed in 4% paraformaldehyde for 10 minutes and diluted to 1% in PBS. Cells were permeabilized in 0.5% Triton-X for 10 minutes and stained with 1:300 KC57-FITC (Beckman Coulter 6604665). Cells were read on a BD FACSCANTO II (Fred Hutch Flow Cytometry Core) and analyzed in FlowJo. For cell surface marker staining, cells were washed twice in PBS, stained in PBS/1% BSA, incubated at 4°C for 1 hr, washed twice in PBS, and analyzed on the Canto 2 flow cytometer (Fred Hutch Flow Cytometry Core).

### Western Blotting

Cells were lysed in 2X SDS-SB lysis buffer (10% glycerol, 2% BME, 6% SDS, 62.5 mM Tris-HCl pH 6.8), boiled at 95*C, sonicated for one minute and resolved by NuPAGE 4–12% Bis-Tris Gel (Invitrogen). Following transfer to a PVDF membrane and blocking in PBS/5%milk for 1 hour, blots were probed with antibodies for 1 hour or overnight, washed in PBST, probed with HRP secondary, washed in PBST and bands visualized with SuperSignal West Femto Maximum Sensitivity Substrate (ThermoFisher #34095). Blots were visualized on a BioRad Chemidoc MP.

### Viral infectivity assays

Cells were pre-stimulated with IFNα 24 hours prior to infection. Virus and 20μg/mL DEAE-Dextran in RPMI were added to cells, spinoculated for 20 minutes at 1100xg, and incubated overnight at 37°C. Cells were washed the next day and re-suspended in RPMI supplemented with IFNα. For experiments to assay ISGs affecting late steps in viral replication, cells were spinoculated at 1100x*g* for 20 minutes with HIV-1_LAI_ or Vpu-deficient HIV-1_LAI_ (HIV_LAI_Δ*vpu*) at an MOI of 0.4, incubated at 37°C for 16 hours, and then treated with 1000 mU/mL IFNα. 24 hours post infection, cells were washed of virus and re-suspended in interferon containing media with 1μg/mL T-20 entry inhibitor (NIH AIDS Reagent Program, Division of AIDS, NIAID, NIH: Enfuvirtide #9409).

### Virus release (p24 ELISA and RT assay)

p24 ELISA on cell culture supernatants was performed with a HIV-1 p24 Ag Assay (Coulter). Reverse transcriptase activity in viral supernatants was measured using the RT activity assay as described (Roesch et al., 2018; Vermeire et al., 2012). A stock of HIV-1_LAI_ virus was titered multiple times, aliquoted at −80°C and used as the standard curve in all assays.

## SUPPLEMENTAL FIGURES

**Supplemental Figure S1.**
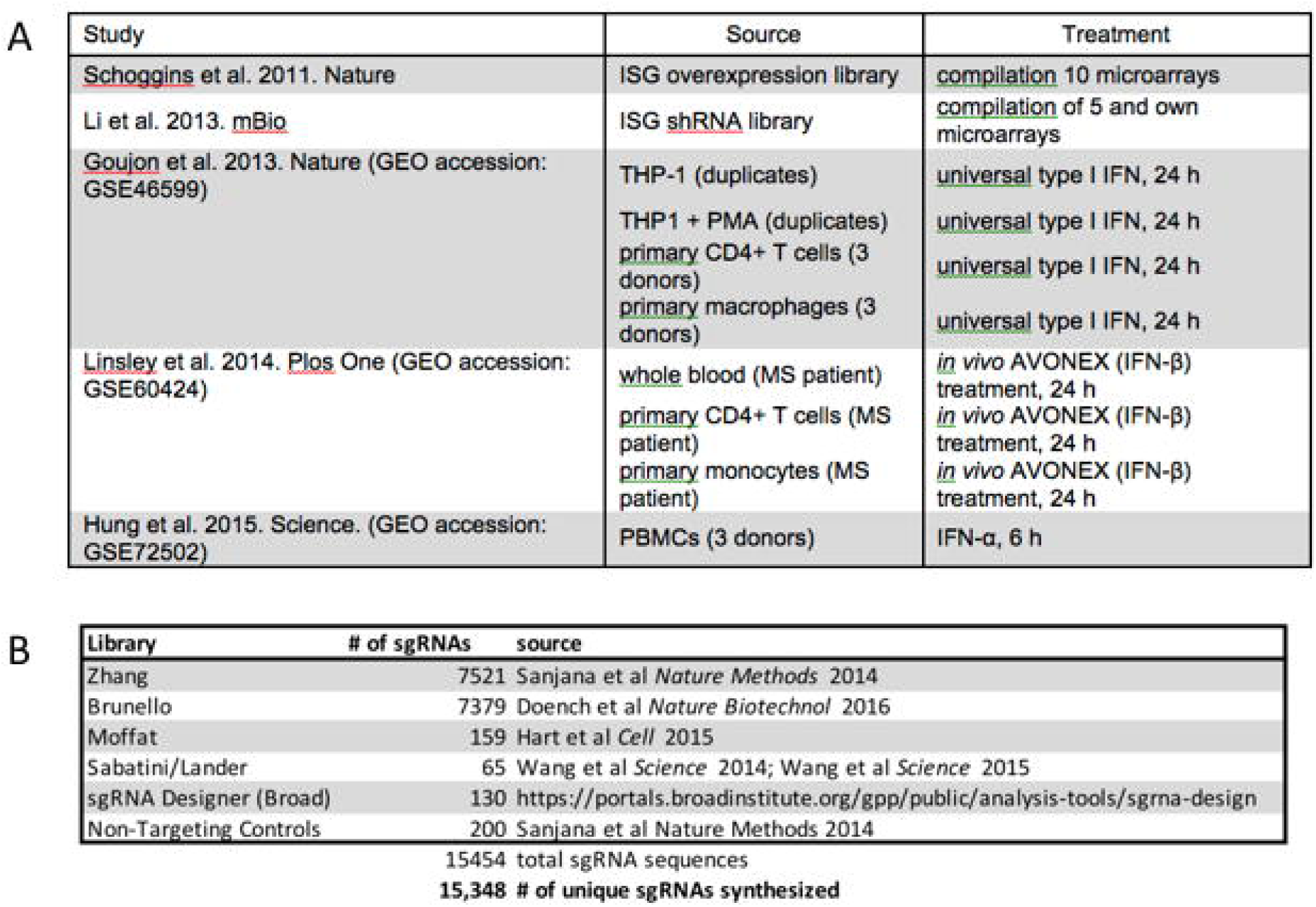
PIKA_HIV_ ISG library gene and sgRNA selection. <u>A</u>: ISG study, source and treatment conditions for ISGs selected for inclusion in the PIKA_HIV_ library. <u>B</u>: Sources for sgRNA sequences included in the PIKA_HIV_ library.

**Supplemental Figure S2.**
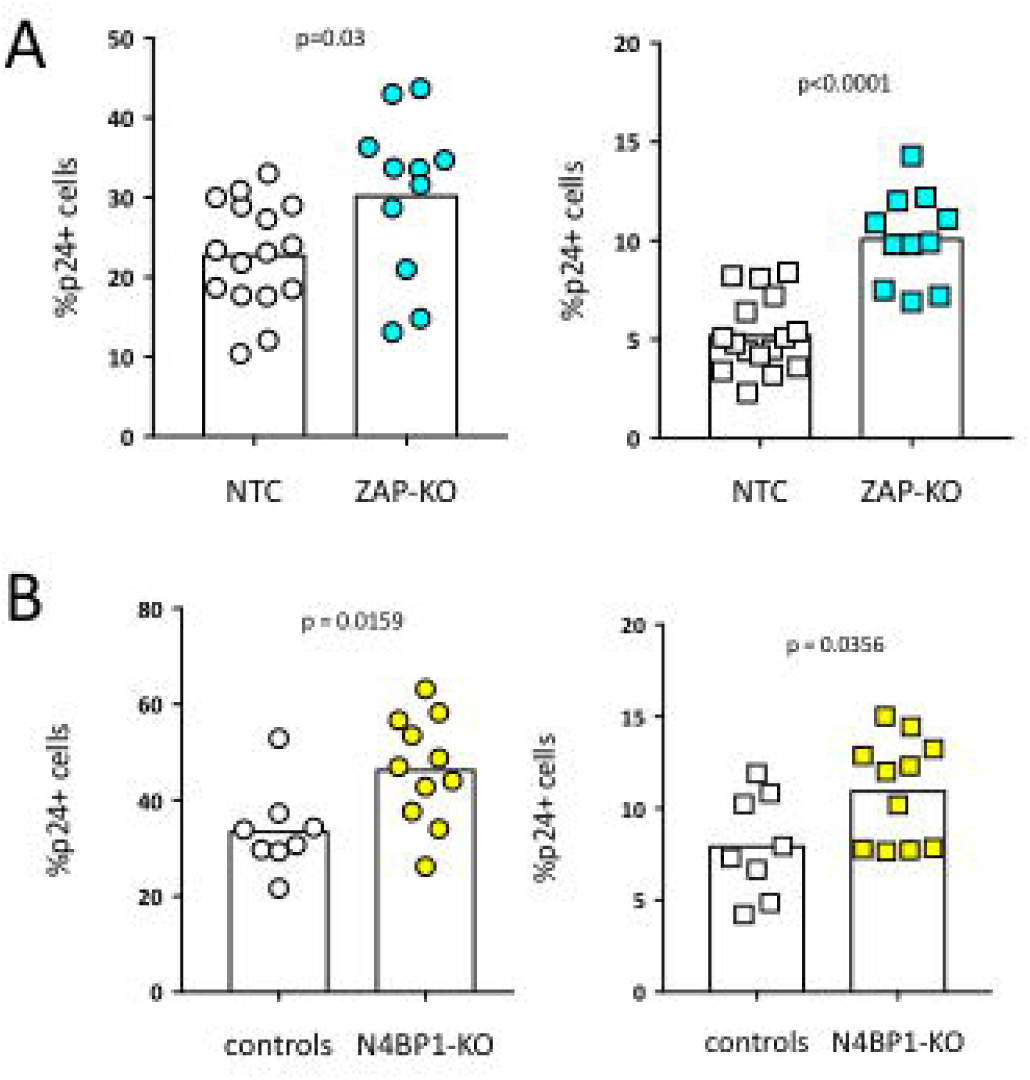
ZAP and N4BP1 are modest inhibitors of HIV infection. <u>A</u>: Control (gray) and ZAP-KO (cyan) clonal THP-1 cell lines were infected and %p24+ cells was measured 2 days post-infection. <u>Left</u>: no IFN. <u>Right</u>: pretreatment with 200U/mL IFNα. <u>B</u>: Control (gray) and N4BP1-KO (yellow) clonal THP-1 cell lines were infected and %p24+ cells was measured 2 days post-infection. <u>Left</u>: no IFN. <u>Right</u>: pretreatment with 200U/mL IFNα.

## SUPPLEMENTAL TABLES

**Supplemental Table S1. ISG Gene Targets in PIKA_HIV_ library.**

**Supplemental Table S2. 20bp sgRNA sequences in the PIKA_HIV_ library.**

**Supplemental Table S3. Log_2_FC analysis of sgRNA enrichment (vRNA:gDNA) for wt THP-1 cells pretreated with IFNα.**

**Supplemental Table S4. IFN Differential Expression Analysis in THP-1 cells and individual and combined MAGeCK Gene Analysis for PIKA_HIV_ screen in ZAP-KO THP-1 cells pretreated with IFNα.**

**Supplemental Table S5. Oligos and primers.**

**Supplemental Table S6. KO scores as determined by ICE analysis for each KO pool. The r^2^ value for each reflects the confidence in the predicted KO score.**

